# p53 dosage impedes Kras^G12D^- and Kras^Q61R^-mediated tumorigenesis

**DOI:** 10.1101/2023.09.17.558140

**Authors:** Özgün Le Roux, Jeffery I. Everitt, Christopher M. Counter

## Abstract

Mice engineered with G12D versus Q61R mutant of Kras exhibit differences in the number and grade of tumors. Namely, the incidence or grade of oral or forestomach squamous epithelial lesions was more prevalent in the Kras^G12D^ background while hematolymphopoietic disease was more prevalent in the Kras^Q61R^ background. Loss of the *Trp53* gene encoding p53 enhances the ability of oncogenic Kras to initiate tumorigenesis in carcinogen and genetic models of lung cancer, while an extra copy of *Trp53* (*Super p53*) was recently shown to suppress Kras-induced tumorigenesis in a genetic model of this disease. Given this, we evaluated whether such an extra copy of *Trp53* would alter tumorigenesis upon global activation of a modified *Kras* allele engineered with either a G12D or Q61R mutation. We report that an increase in p53 dosage generally reduced tumor number or grade across a number of organs in a manner largely independent of the type of Kras mutation, which was sufficient to extend lifespan in the less aggressive background of a *Kras^G12D^* initiating mutation.

## Introduction

The RAS superfamily of small GTPases is comprised of the genes *KRAS*, *NRAS*, and *HRAS* in humans (*Kras*, *Nras*, and *Hras* in mice) [1]. Activated growth factor receptors are known to lead to the switching of RAS from an inactive GDP-bound to an active GTP-bound state. Once activated, RAS recruits proteins with RAS-binding domains or Ras-associating domains, thereby activating these proteins to propagate signaling. Hydrolysis of GTP returns RAS to the inactive GDP-bound state, terminating the signaling [2]. However, in a fifth or more of human cancers, point mutations at one of the three hotspot sites render RAS constitutively active, which is well established to initiate tumorigenesis [1-4]. Between the three RAS genes, three hotspots, with six possible substitutions there are more than 50 potential oncogenic mutations, yet cancers tend to have a particular bias, or a ‘RAS mutation tropism’, towards an often unique subset of these mutations [5]. Case in point, the most common RAS mutations in lung cancer are G12C/V/D in KRAS [6], but Q61R/K in NRAS in melanoma [7].

Expression of different RAS mutants in mice can also affect the nature of tumors arising. For example, expressing a *Kras^G12D^*, but not an *Nras^G12D^* allele in the skin of mice leads to melanoma [8]. Three oncogenic *Kras* mutations were more commonly detected in tumors arising in mice infected with an sgRNA library designed to generate all 12 oncogenic mutations at codons G12 and G13 [9]. We recently created Cre-inducible (lox-STOP-lox or LSL) *Kras* alleles encoded by common (com) codons and either of the two biochemically distinct mutations G12D or Q61R. Globally activating these two via Cre expressed from the ubiquitous *Rosa26* locus led to differences in the severity and in some cases even type of tumors arising in a manner specific to mutation type. Namely, the incidence or grade of oral or forestomach squamous epithelial lesions was more prevalent in the Kras^G12D^ background while hematolymphopoietic disease was more prevalent in the Kras^Q61R^ background [10].

The patterns of tumors induced by these two alleles is ostensibly a product of how a normal cell responds to different activation levels or specific mutant oncoproteins. In this regard, oncogenic RAS can both induce proliferation or senescence, the latter of which is mediated in part by the tumor suppressor p53 [11]. Recently, “*super p53*” mice encoding an extra copy of *Trp53* [12] were shown to suppress spontaneous Kras-driven lung and lymphoma tumorigenesis arising in an *LA1-Kras^G12D^* background, but not radiation-induced lymphomas [13]. This prompted us to ask whether an increase in p53 dosage would alter the severity or type of tumors arising between different oncogenic mutations. We thus generated mice with the two experimental genotypes of *Kras^LSL-comG12D/+^*;*Super p53* and *Kras^LSL-comQ61R/+^*;*Super p53* and the two control genotypes of *Kras^LSL-comG12D/+^* and *Kras^LSL-comQ61R/+^* in the *Rosa26^CreERT2/+^* background, which allows ubiquitous Cre expression in a broad range of tissues [14]. The *Kras^LSL^* alleles were activated by tamoxifen treatment and the mice were analyzed for the presence of hematolymphopoietic, oral and forestomach, and lung tumors, as well as tumors in other organs. We report here that an increase in *p53* dosage generally suppressed tumor number or type across several organs, largely independent of the type of Kras mutation, which was sufficient to extend lifespan in the less aggressive *Kras^G12D^* background. This implies that regardless of whether a tissue is preferentially induced by the level or type of mutant of Kras, an extra copy of *Trp53* may suppress initiation and/or progression of the resulting tumors.

## Results

### The experimental design to explore the influence of p53 dosage on tumorigenesis induced by different Kras mutations

Mice with *Rosa26^CreERT2/CreERT2^* alleles were crossed with those having either of a *Kras^LSL-comG12D/+^* or *Kras^LSL-comQ61R/+^* genotype twice to create *Rosa26^CreERT2/CreERT2^;Kras^LSL-comG12D/+^* and *Rosa26^CreERT2/CreERT2^*;*Kras^LSL-comQ61R/+^*mice. These mice were then crossed with *Super p53* mice to create *Kras^LSL-comG12D/+^*;*Super p53* and *Kras^LSL-comQ61R/+^*;*Super p53* mice in the *Rosa26^CreERT2/+^*background. Admittedly this breeding strategy yielded significantly small number of mice (around 5%) having a *Kras^LSL^* background without an extra allele of *Trp53* therefore all tumorigenesis comparisons were made against the original tumorigenesis study in *Rosa26^CreERT2/CreERT2^*;*Kras^LSL-comG12D/+^* and *Rosa26^CreERT2/CreERT2^*;*Kras^LSL-comQ61R/+^* mice that lacked the extra allele of *Trp53* [10]. We thus generated experimental cohorts of 5 to 9 mice in *Kras^LSL-comG12D/+^*;*Super p53* and *Kras^LSL-comQ61R/+^*;*Super p53* genotypes **(S1 Fig)**, which were compared to the control cohorts of 23 to 24 mice in *Kras^LSL-comG12D/+^* or *Kras^LSL-comQ61R/+^*genotypes in the *Rosa26^CreERT2/+^* background as previously reported [10]. All mice were injected with tamoxifen, which we previously extensively validated to result in recombination (and thus activation) of both the *Kras^LSL-comG12D/+^* and *Kras^LSL-comQ61R/+^* alleles in many different organs [10]. Mice were regularly monitored until moribundity, at which time they were humanely euthanized **(Fig 1A)**.

**Fig 1.**
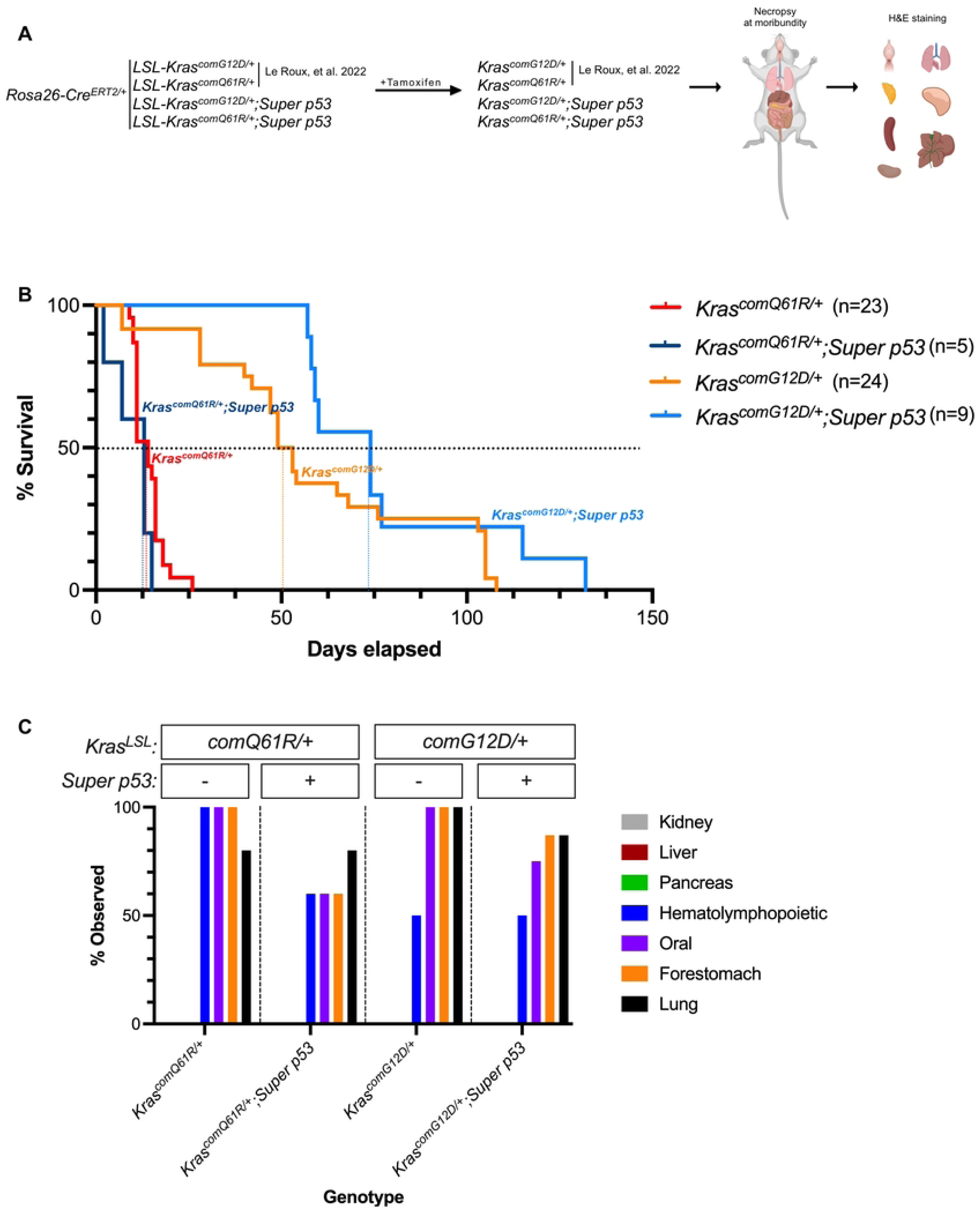
The effect of p53 dosage on Kras-driven tumorigenesis. **(A)** Experimental design to study the effect of the *Super p53* allele on tumorigenesis upon globally activating *Kras^LSL-comG12D^* versus *Kras^LSL-comQ61R^* alleles by tamoxifen in a *Rosa26^CreER2/+^* background. **(B)** Kaplan-Meier survival curve of *Rosa26^CreER2/+^* mice with the indicated genotypes (0 days refers to the last tamoxifen injection). Dotted lines: 50% survival. Pairwise comparison for statistical analysis is provided in **S2 Fig**. **(C)** Percent of *Rosa26^CreER2/+^* mice of the indicated genotypes (*n*=5-8) with the described tumor types. Examples of H&E-stained slides of the indicated tissues are provided in **S3 Fig.**

### An increase in p53 dosage provides an overall survival benefit in mice in which Kras^G12D^ is globally activated

The time of moribundity, which was previously shown to precede death due to cancer [10], was plotted for all four cohorts using the Kaplan-Meier method [15]. We previously reported the median survival of *Kras^comG12D/+^* and *Kras^comQ61R/+^* mice after tamoxifen injections was 14 and 51 days, respectively, in the *Rosa26^CreERT2/+^*background [10]. With increased p53 dosage, the median survival was unchanged in the *Kras^comQ61R/+^*cohort but increased by 23 days (51 to 74 days) in the *Kras^comG12D/+^*mice, which equates to 45% increase in survival **(Fig 1B)**. Although pairwise comparisons did not show statistically significant differences between experimental and control cohorts using the Log rank test **(S2A,B Fig)**, there was a statistically significant increase (*p* value <0.05) in the median survival of the *Kras^comG12D/+^* mice with an extra allele of *Trp53* using the Gehan-Breslow-Wilcoxon test **(S2B Fig)**, which weighs early time points higher [16]. Thus, an extra copy of *p53* appears to extend lifespan in the context of the less aggressive *Kras^comG12D/+^*allele.

### An increase in p53 dosage alters the severity of tumorigenesis across multiple organs

To examine the effect of an increase in p53 dosage on tumorigenesis, the lung, thymus, stomach, pancreas, spleen, oral epithelia, liver, and kidney were removed from five to eight mice from each of the four cohorts and paraffin embedded and H&E stained **(Fig 1A)**. Two slides from each of these eight organs from all animals in the study were analyzed for the incidence and grading of tumors by a board-certified veterinary pathologist **(S3 Fig)**. In terms of the number of mice with tumors in these organs, we find, not surprisingly, that organs previously shown to be resistant to Kras-induced tumorigenesis [10], namely the kidney, liver, and pancreas, remained tumor-free **(Fig 1C)**. However, with two exceptions, there was a reduction in mice with oral, forestomach, lung, and hematolymphopoietic lesions in the both the *Kras^comG12D/+^*and *Kras^comQ61R/+^* mice in the *Super p53* background **(Fig 1C)**. The two exceptions were the number of animals with hematolymphopoietic and lung lesions was the same in *Kras^comG12D/+^*and *Kras^comQ61R/+^* mice, respectively, with versus without an extra copy of *Trp53* **(Fig 1C)**. Whether this reflects underlying biology or the limited number of mice with an extra copy of *p53* remains to be determined. Thus, the additional copy of *Trp53* generally reduced the incidence of tumors detected in a wide spectrum of organs in a manner largely independent of mutation type.

### Oral and forestomach squamous epithelial lesions

We previously reported that oral and forestomach squamous epithelial lesions were more prevalent and aggressive in a *Kras^LSL-comG12D/+^* compared to a *Kras^LSL-comQ61R/+^* genotype, suggesting that these two tissues were particularly sensitive to the ability of the G12D mutant of Kras to induce tumorigenesis [10]. To explore the impact of an extra allele of *Trp53* on the tumorigenesis in these two tissues, the aforementioned H&E-stained slides were analyzed and graded as either mild, moderate, or marked atypical squamous hyperplasia (ASH) or as squamous papilloma. In both the *Kras^comG12D/+^*and *Kras^comQ61R/+^* background, increased p53 dosage reduced the severity of the oral squamous epithelial lesions **(Fig 2A)**. Namely, the percentage of animals with oral squamous epithelial lesions decreased with increased p53 dosage in the *Kras^comG12D/+^* and *Kras^comQ61R/+^* background by 20% and 40%, respectively. Furthermore, the percentage of mice with the *Kras^comG12D/+^* alleles that had the most severely graded oral lesions of squamous papilloma decreased from 100% to 80% with an extra copy of *Trp53*, while there was a clear shift from more severe lesions of marked and moderate ASH to less severe lesions of mild ASH or no lesions in the *Kras^comQ61R/+^* background with increased p53 dosage **(Fig 2A)**. The effect was similar in forestomach squamous epithelial lesions, although less pronounced **(Fig 2B)**. Specifically, the percentage of animals with forestomach squamous epithelial lesions decreased with increased p53 dosage in *Kras^comG12D/+^* and *Kras^comQ61R/+^* background by 25% and 40%, respectively. Furthermore, in both *Kras^comG12D/+^* and *Kras^comQ61R/+^*background, increasing p53 dosage led to a shift from more severe to less severe squamous epithelial lesion types in the forestomach, which was more noticeable in the *Kras^comQ61R/+^* genotype **(Fig 2B)**. The observation that the number and grade of oral and forestomach squamous epithelial lesions decreased in the *super p53* background supports the contention that an additional copy of *Trp53* suppresses both tumor initiation and progression independent of the nature of the initiating Kras mutation.

**Fig 2.**
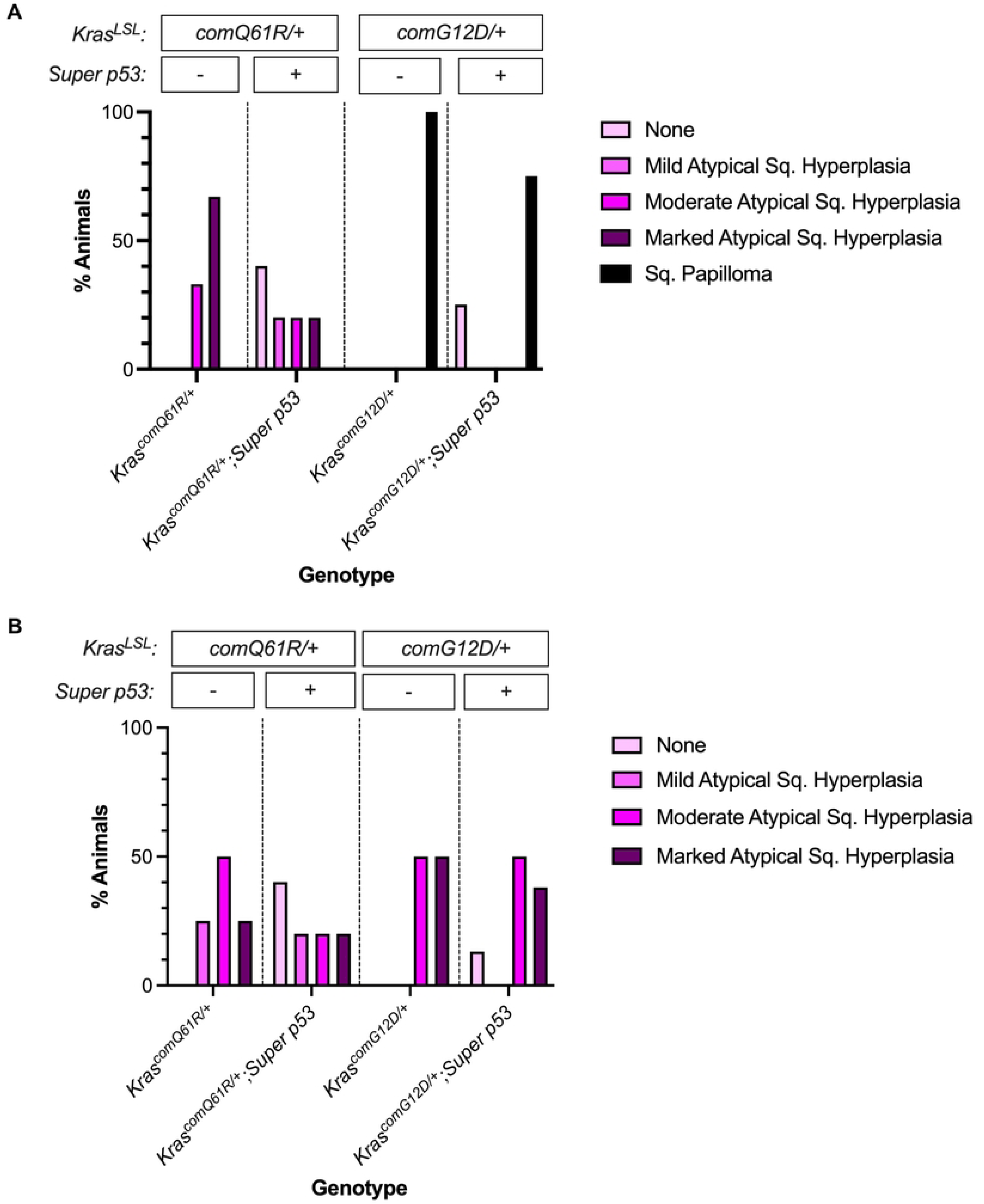
The effect of p53 dosage on Kras-driven oral and forestomach tumorigenesis. Percent of *Rosa26^CreER2/+^* mice of the indicated genotypes (*n*=5-8) with the indicated grades of **(A)** oral and **(B)** forestomach squamous epithelial lesions at moribundity endpoint.

### Hematolymphopoietic lesions

We previously reported that hematolymphopoietic disease was more severe either through increased Kras expression or in the *Kras^comQ61R/+^* mutant background, suggesting that this tissue was particularly sensitive to the level of active Kras to induce tumorigenesis [10]. Given this, we compared the number of mice with myeloproliferative disease as well as the number and grade of mice with lymphomas. Specifically, the aforementioned H&E slides of all tissues from each of the four genotypes were analyzed for the presence of hematolymphopoietic lesions. As previously reported, the myeloproliferative disease was only detected in mice with Q61R mutation [10] and the percentage of animals with myeloproliferative disease decreased from 100% to 60% with an extra copy of *Trp53* in this background **(Fig 3A)**. Consistent with decreased incidence, percentage of mice having the *Kras^comQ61R/+^* genotype with myeloproliferative infiltrates in spleen, kidney, liver, pancreas, and lung was decreased with increased p53 dosage **(S4A Fig)**. When we compared the impact of extra *Trp53* allele on G12D-induced hematolymphopoietic lesions, we observed that the percentage of animals with lymphoma did not change **(Fig 3A)**, although the distribution of lymphoma infiltrates varied **(S4B Fig)** as well as perhaps also the type of lymphomas **(Fig 3B)**, namely a shift from polymorphic cytology to a thymic lymphoblastic T-cell (CD3+) malignant lymphoma **(S4C Fig** and not shown). Thus, an additional copy of *Trp53* appears to suppress tumor initiation and/or progression in the hematolymphopoietic compartment, particularly in the *Kras^comQ61R/+^*background.

**Fig 3.**
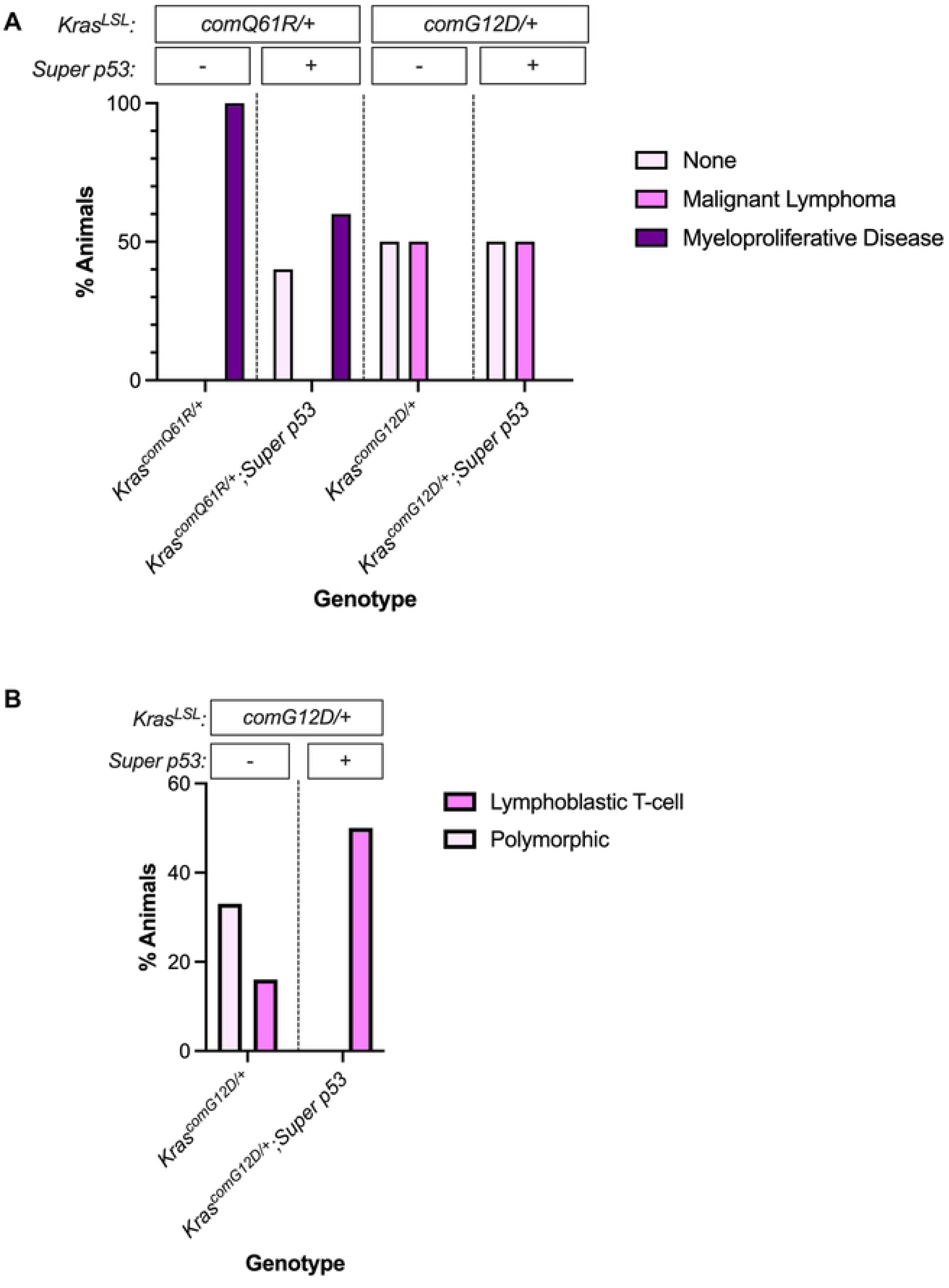
The effect of p53 dosage on Kras-driven hematolymphopoietic tumorigenesis. Percent of *Rosa26^CreER2/+^* mice of the indicated genotypes (*n*=5-8) with the indicated type of **(A)** hematolymphopoietic disease and **(B)** lymphomas.

### Lung lesions

Upon comparing the incidence and grading of the lesions in the lungs of all four cohorts, we found that with increased p53 dosage, the severity of the peripheral lesions was reduced in both *Kras^comG12D/+^* and *Kras^comQ61R/+^* mice. Specifically, with extra copy of *Trp53*, percentage of animals with peripheral hyperplasia decreased from 100% to 87% in the *Kras^comG12D/+^* background, while peripheral hyperplasia were actually absent in the *Kras^comQ61R/+^* background **(Fig 4A)**. However, there was no difference in other types of lung tumors (bronchiolar hyperplasia and adenomas), other than an increase in bronchiolar hyperplasia in the *Kras^comG12D/+^* background **(S5A,B Fig)**. Thus, the addition of an extra copy of *Trp53* appeared to have the greatest effect on peripheral lung hyperplasia.

**Fig 4.**
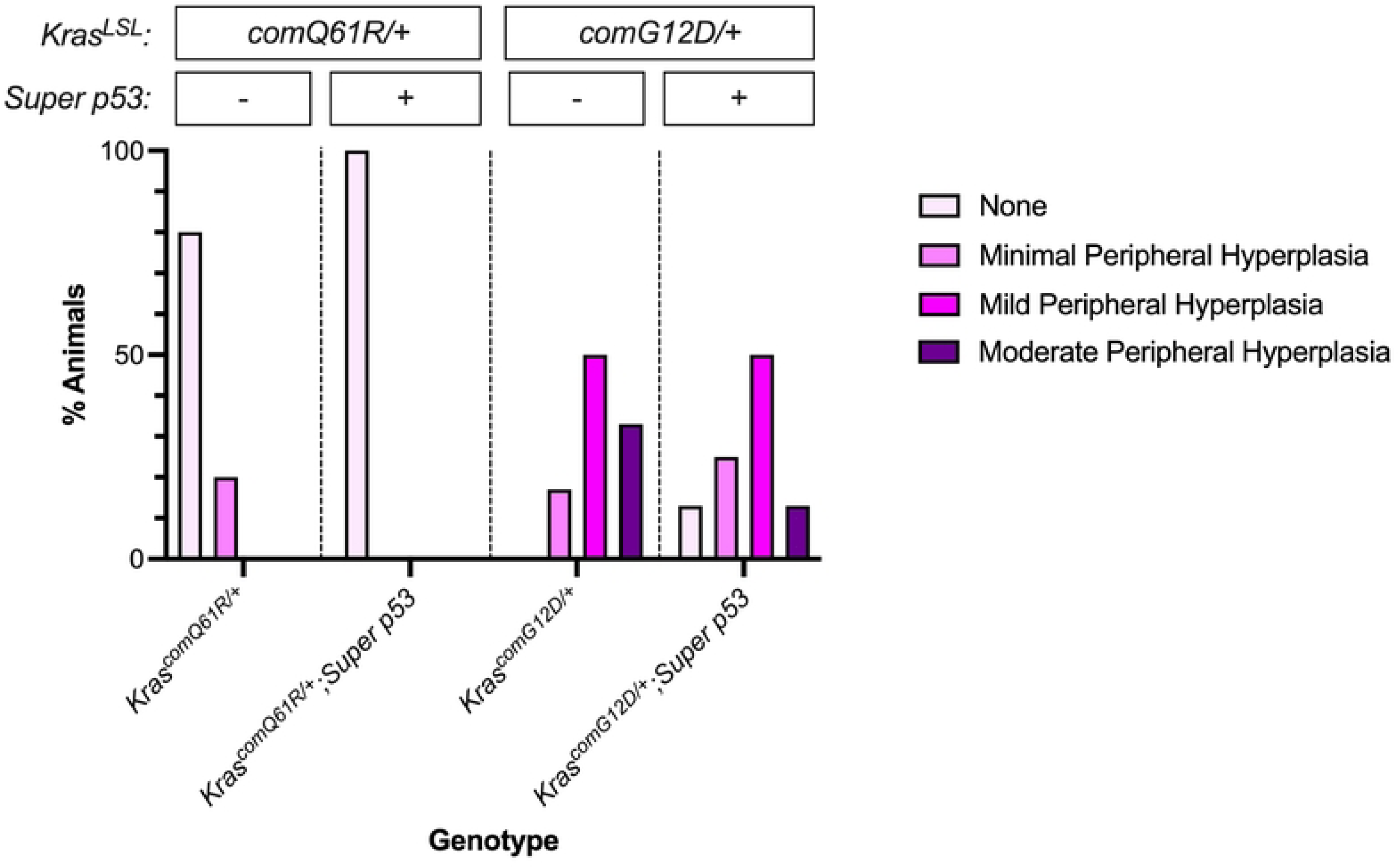
The effect of p53 dosage on Kras-driven lung tumorigenesis. Percent of *Rosa26^CreER2/+^*mice of the indicated genotypes (*n*=5-8) with the indicated grades of the peripheral lung hyperplasia.

## Discussion

We report that increasing p53 dosage extended lifespan in *Rosa26^CreERT2/+^;Kras^LSL-comG12D/+^*mice by one statistical comparison, and this was associated with a decrease in the incidence and/or grade of lesions in the four organs pathologically analyzed, namely the oral cavity, forestomach, hematolymphopoietic compartment, and lung (except for lung bronchiolar hyperplasia). An additional copy of *Trp53* did not, however, extend lifespan in the *Rosa26^CreERT2/+^;Kras^LSL-comQ61R/+^* mice. Nevertheless, the number of these mice without lesions in the oral cavity, forestomach, and hematolymphopoietic compartment increased, and peripheral hyperplasia in the lung were entirely absent. While the similar lifespan between *Kras^LSL-comQ61R/+^* and *Kras^LSL-comQ61R/+^*;*Super p53* mice may reflect a mutation-specific effect, we favor a model whereby the median survival of 2 weeks is simply too short for an extra copy of *Trp53* to alter lifespan. We also note that the effect of an extra copy of *Trp53* did not always suppress tumorigenesis. For example, the percentage of animals with lung tumors in the *Kras^comQ61R/+^* genotype or hematolymphopoietic cancers in the *Kras^comG12D/+^*genotype did not change with increased p53 dosage **(Fig 1C,3A)**. Similarly, *Kras^comG12D/+^* induced similar number of adenomas per animal with and without the extra allele of *Trp53* **(S5B Fig)**. We note the following caveats to this study. First, due to the breeding strategy, the number of littermates with the control genotypes *Rosa26^CreERT2/+^*;*Kras^LSL-comG12D/+^* or *Rosa26^CreERT2/+^*;*Kras^LSL-comQ61R/+^*was extremely limited, and thus we chose to make comparisons to the control genotypes reported in our previous study as all experimental steps were performed in the same manner. Thus, some trends may be the product of the natural variation in tumorigenesis rather than a bona fide phenotypic difference. However, the reduction in the number and/or grade of tumors was a reoccurring theme in the *Super p53* background, supporting this as a real phenomenon. Second, we have not measured the level of p53 protein in the *Trp53^+/+^* versus *Super p53* backgrounds to confirm an increase in the latter is linked to the observed phenotypes. However, by the very nature of providing an extra copy of this gene, p53 protein was previously demonstrated to increase with genotoxic insult in mouse embryonic fibroblasts with subsequent decrease in cell cyle arrest [12]. Third, we appreciate that the *Kras^LSL-comG12D/+^* and *Kras^LSL-comQ61R/+^* alleles have altered the normal structure of the *Kras* gene, and hence limit the degree that these two alleles can be compared to other modified *Kras* alleles. Nevertheless, by using two alleles constructed in the identical fashion, meaningful comparisons could be made. In summary, with these exceptions and caveats, the extra copy of *Trp53* appears to be generally suppressive regardless of the type of oncogenic Kras mutation used to initiate tumorigenesis, although the effect was clearly greater when tumorigenesis is less aggressive. While admittedly very speculative, we suggest that given these findings, perhaps then naturally occurring variations in the level of p53 protein in the human population may influence the likelihood that a spontaneously arising KRAS mutation initiates tumorigenesis, which may find value in predicting susceptibility to Kras-driven cancers or in development of therapeutic strategies against KRAS-induced cancers.

## Materials and methods

### Mouse strains

*Kras^LSL-comG12D/+^* and *Kras^LSL-comQ61R/+^*mice were previously described and were from a pure 129 background [10]. *Super p53* mice were kindly provided by David Kirsch (University of Toronto) and were from C57BL/6 background. *Rosa26^CreERT2/CreERT2^*mice were obtained from The Jackson Laboratory (strain #008463). All animals derived in subsequent crosses were selected by genotyping using the genotyping methods previously described [10, 12]. All animal experiments were approved by Duke IACUC.

### Tumorigenesis studies

*Kras^LSL-comG12D/+^* and *Kras^LSL-comQ61R/+^*mice were crossed with mice with *Rosa26^CreERT2/CreERT2^* allele (Jackson Laboratory, strain 008463) to generate *Rosa26^CreERT2/+^*;*Kras^LSL-comG12D/+^*and *Rosa26^CreERT2/+^*;*Kras^LSL-comQ61R/+^* mice. These *Kras^LSL^* mice in the *Rosa26^CreERT2/+^* background were then crossed with the *Rosa26^CreERT2/CreERT2^*mice again to generate *Rosa26^CreERT2/CreERT2^*;*Kras^LSL-comG12D/+^* and *Rosa26^CreERT2/CreERT2^*;*Kras^LSL-comQ61R/+^*mice. *Kras^LSL^* mice with homozygous Cre alleles were then crossed with the *Super p53* mice to generate the littermates of *Kras^LSL-comG12D/+^* and *Kras^LSL-comG12D/+^*;*Super p53*, and *Kras^LSL-comQ61R/+^* and *Kras^LSL-comQ61R/+^*;*Super p53* mice in the *Rosa26^CreERT2/+^* background. This crossing strategy generated *Kras^+/+^;Super p53* mice at the ratio of 80%, leading to relatively small number of *super p53* mice harboring a *Kras^LSL^* allele in the *Rosa26^CreERT2/+^*background (around 10%), while the ratio of littermates with the *Kras^LSL^*allele in the *Rosa26^CreERT2/+^* background without the extra *Trp53* was around 5% **(S1 Fig)**. Despite our efforts to compare littermates of *Kras^LSL^*mice with and without an extra allele of *Trp53,* with all littermates in the *Rosa26^CreERT2/+^* background, this strategy came at the cost of yielding significantly small number of control littermates with a *Kras^LSL^* background without an extra allele of *Trp53* after multiple rounds of crosses. Using this strategy, we generated experimental cohorts of 7 to 9 mice with random distribution of males and females in *Kras^LSL-comG12D/+^*;*Super p53* and *Kras^LSL-comQ61R/+^*;*Super p53* genotypes. For comparison of median survival, comparisons were made against the control cohorts we reported previously with 23 to 24 mice with random distribution of males and females in *Kras^LSL-comG12D/+^*or *Kras^LSL-comQ61R/+^* genotypes in the *Rosa26^CreERT2/+^* background [10]. For comparison of tumorigenesis, we chose to limit the number of control animals in the study to a similar number in the experimental cohort to make statistical power evenly distributed across cohorts. Thus, tumorigenesis comparisons were made against 5 to 6 randomly chosen mice from the control cohorts with random distribution of males and females in *Kras^LSL-comQ61R/+^* and *Kras^LSL-comG12D/+^*genotypes in the *Rosa26^CreERT2/+^* background[10]. Tumorigenesis studies were performed in the same manner as reported previously **(Fig 1A)** [10]. Namely, tamoxifen (Sigma-Aldrich, T5648-5G, CAS# 10540-29-1) was dissolved in corn oil (Sigma-Aldrich, C8267) and filter sterilized. At six to eight weeks of age, experimental cohorts of 7 to 9 mice with random distribution of males and females in *Kras^LSL-comG12D/+^*;*Super p53* and *Kras^LSL-comQ61R/+^*;*Super p53* genotypes were injected intraperitoneally with 250 μg/g body weight of tamoxifen four times every 48 hrs. Tamoxifen injections were performed under anestesia by isoflurane to prevent suffering. After recombination of the *LSL* cassette with tamoxifen, mice were observed daily during injections, 1 week after last injection, and weekly thereafter. Mice were humanely euthanized with carbon dioxide inhalation followed by removal of vital organs within 2 hrs of detection of moribundity humane endpoint. To prevent suffering and pain, we defined moribundity humane endpoint as any of the visible signs of sudden behaviroal change, poor/hunched posture, lost hair coat condition, sudden activity level change, painful facial expression, signs of pain that was not anticipated by the study plan, weight loss of exceeding 15% compared to an age-matched reference, and cardiopulmonary disorders. Selected tissues, including lung, thymus, stomach, pancreas, spleen, oral epithelia, liver, and kidney, were removed at necropsy followed by fixation in 10% formalin (VWR, 89370-094) for 24-48 hrs, then were stored in 70% ethanol (VWR, 89125-166) until routine processing. 2 animals from the *Kras^LSL-comQ61R/+^*;*Super p53* genotype were found dead within 24 hrs of the first tamoxifen injection, thus, were not included in either of the reported survival or tumorigenesis studies as we could not perform necropsy in a timely manner. Tissues were sliced with one-to-two-millimeter thickness. Tissue slices were embedded in paraffin with the flat sides down, sectioned at a depth of 5 μm, and stained by the H&E method by IDEXX Laboratories. All H&E slides were evaluated by a board-certified veterinary pathologist with experience in murine pathology in a blinded fashion.

### Statistical analysis

Statistical analyses were performed using GraphPad Prism software version 9.5.1 (GraphPad Software). Pairwise comparisons of the survival plots of each cohort with and without an extra allele of *Trp53* were performed with Log-rank (Mantel-Cox) and Gehan-Breslaw-Wilcoxon test[17] **(S2B-C Fig)**. For comparisons of number of lesions per animal, one-way ANOVA with Bonferroni’s multiple-comparisons test with a single pooled variance and a 95% CI were used **(S5A-B Fig)**. A *p*-value of less than 0.05 was considered statistically significant.

## Acknowledgements

We thank members of the Counter Laboratory for technical assistance, and David Kirsch for advice. All mouse and organ pictures were created with Biorender.com. This work was supported by the National Cancer Institute (R01CA269272 to CMC) and the Shared Resources of the Duke Cancer Institute (P30CA014236). The authors declare no competing interests.

## Author contributions

Özgün Le Roux: Conceptualization, Resources, Data curation, Formal analysis, Validation, Investigation, Visualization, Methodology, Writing – original draft, Writing – review and editing Jeffery I. Everitt: Data curation, Methodology, Writing – review and editing Christopher M. Counter: Conceptualization, Resources, Supervision, Funding acquisition, Visualization, Writing – original draft, Project administration, Writing – review and editing

## Supporting information

**S1 Fig. Breeding strategy to generate the indicated genotypes.**

**S2 Fig. Pairwise comparisons of survival curves.**

Pairwise comparison of *Rosa26^CreER2/+^* mice upon activating the **(A)** *Kras^LSL-comQ61R^* or **(B)** *Kras^LSL-comG12D^* allele in the absence and presence of the *Super p53* allele from **Fig 1B**. Log-rank and Gehan-Breslow-Wilcoxon test statistical analysis results are shown on the right. *: *p*-value <0.05. ns: not significant. df: degrees of freedom.

**S3 Fig. Examples of H&E-stained tissues.**

Examples of H&E staining of the eight indicated organs removed at moribundity endpoint from *Rosa26^CreER2/+^* mice with the four indicated genotypes. 4X magnification. Scale bar: 600 μm.

**S4 Fig. Hematolymphopoietic infiltrates and types.**

**(A,B)** Percent of *Rosa26^CreER2/+^* mice (*n*=5-8) with hematolymphopoietic infiltrates in the indicated organs upon activating the **(A)** *Kras^LSL-comQ61R^* or **(B)** *Kras^LSL-comG12D^* alleles in the absence and presence of the *Super p53* allele.

**(C)** Examples showing lymphoma infiltrates in an H&E-stained section of the indicated organs from a *Rosa26^CreER2/+^;Kras^LSL-comG12D^*versus *Rosa26^CreER2/+^;Kras^LSL-comG12D^;Super p53* mouse.

**S5 Fig. The effect of p53 dosage on the types of Kras-driven lung tumorigenesis.**

Number of lung **(A)** bronchiolar hyperplasia foci or **(B)** adenomas per animal in *Rosa26^CreER2/+^* mice of the indicated genotypes (*n*=5-8). One-way ANOVA with Bonferroni’s multiple-comparisons test with a single pooled variance and a 95% CI were used. *: mice with extensive myeloid infiltrates that precludes accurate determination of the number of lung lesions. **: *p*<0.001.

## Notes

### Competing Interest Statement

The authors have declared no competing interest.

